# Organization of Neural Population Code in Mouse Visual System

**DOI:** 10.1101/220558

**Authors:** Kathleen Esfahany, Isabel Siergiej, Yuan Zhao, Il Memming Park

**Affiliations:** Ward Melville High School, East Setauket, NY, USA; Department of Computer Science, Cornell University, Ithaca, NY, USA; Department of Neurobiology and Behavior, Stony Brook University, Stony Brook, NY, USA; Institute for Advanced Computational Science, Stony Brook University, Stony Brook, NY, USA

## Abstract

The mammalian visual system consists of several anatomically distinct areas, layers, and cell types. To understand the role of these subpopulations in visual information processing, we analyzed neural signals recorded from excitatory neurons from various anatomical and functional structures. For each of 186 mice, one of six genetically tagged cell-types and one of six visual areas were targeted while the mouse was passively viewing various visual stimuli. We trained linear classifiers to decode one of six visual stimulus categories with distinct spatiotemporal structures from the population neural activity. We found that neurons in both the primary visual cortex and secondary visual areas show varying degrees of stimulus-specific decodability, and neurons in superficial layers tend to be more informative about the stimulus categories. Additional decoding analyses of directional motion were consistent with these findings. We observed synergy in the population code of direction in several visual areas suggesting area-specific organization of information representation across neurons. These differences in decoding capacities shed light on the specialized organization of neural information processing across anatomically distinct subpopulations, and further establish the mouse as a model for understanding visual perception.

## Introduction

Though the mouse has long been neglected as a model for studying neural visual information processing, it has recently emerged as a powerful alternative to primate and other carnivorous species. Mice offer the benefit of large-scale, high-throughput experiments and sophisticated genetic tools for investigating highly specific components of visual perception Arenkiel and Ehlers (2009). However, the use of mice in studying visual perception is currently limited by insufficient knowledge about the functional organization of the mouse visual cortex. Thus, we aim to characterize the population neural code associated with cortical organization of visual information processing.

Visual information is thought to be processed in a series of computations as it travels from the retina to the lateral geniculate nucleus and then through a series of visual cortices Nassi and Callaway (2009). The early visual system processes complex visual stimuli through the simultaneous encoding of different stimulus attributes, such as direction, orientation, and spatial and temporal frequency by individual neurons, while higher order visual cortices process nonlinear features Orban (2008). If we can build a simple population decoder to read out the information made accessible by the neural population (Fig. 1), we can provide insight to which of these features are encoded in specific populations of neurons Graf et al. (2011).

**Figure 1:**
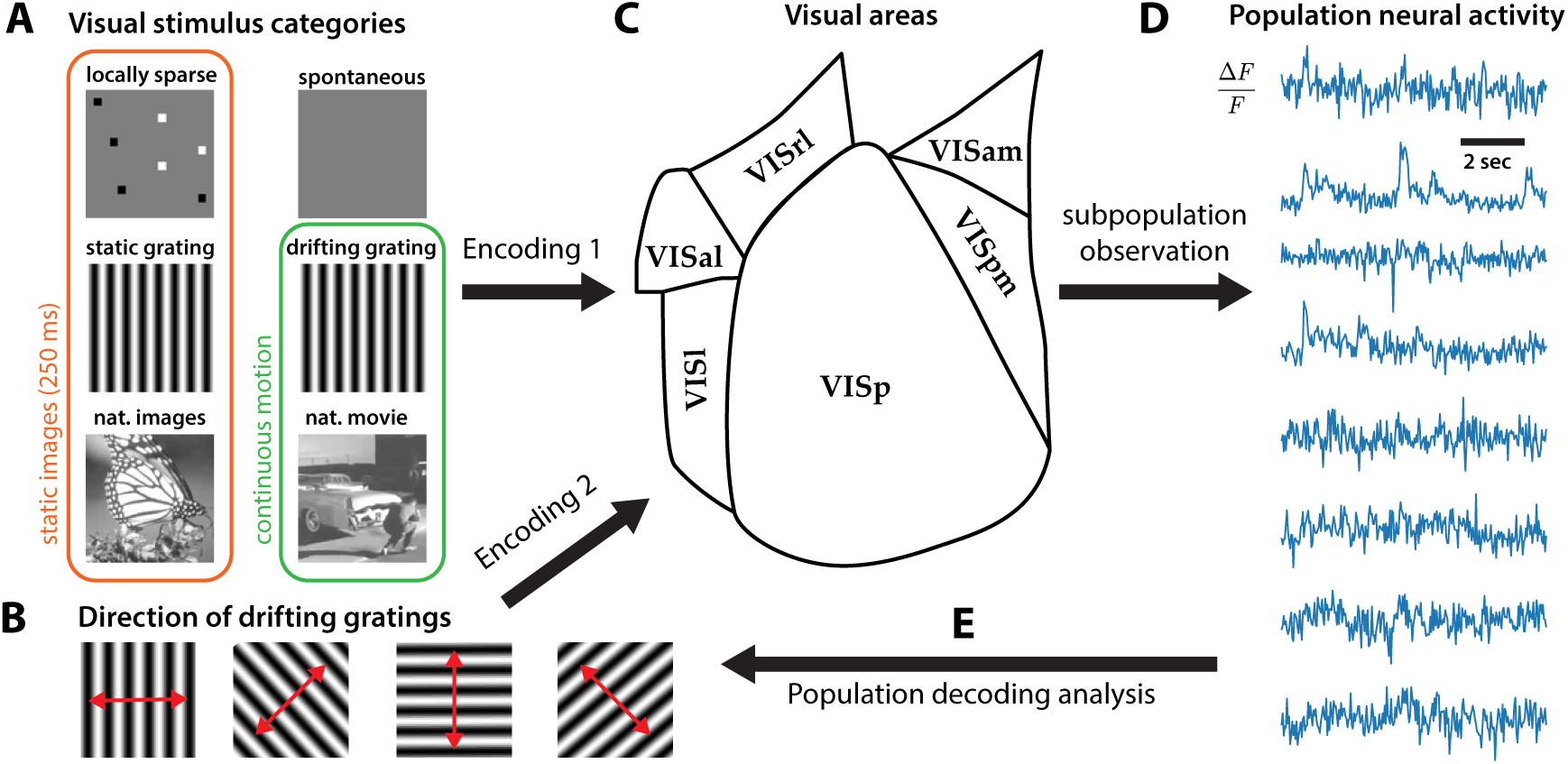
Overview of the population decoding analysis. The neural code of either one of six visual categories (**A**) or one out of eight directions of drifting grating stimulus (**B**) by the excitatory neurons in the mouse visual system (**C**) were analyzed. A specific subpopulation (visual area, cell-type, depth) were targeted and observed while the mice viewed the visual stimuli. From the normalized fluorescence signals from the subpopulation (**D**), we decoded the identity of the stimulus class (**E**). Successful decoding provides evidence for a instantaneous representation of the spatiotemporal signatures of stimuli within the population activity.

The global topographic organization of the mouse visual cortex has been well characterized. Recent studies have retinotopically identified at least ten visual areas with organized and complete representations of the entire visual field Garrett et al. (2014), Marshel et al. (2011), Wang and Burkhalter (2007). However, the neural population code — how information is collectively represented in the neural activity — has remained elusive. While progress has been made in identifying differences between the spatiotemporal information encoded by neurons in different visual areas, prior work has focused on single neurons and lacks analysis at the population level Andermann et al. (2011), Marshel et al. (2011). By decoding neural responses in large neural populations of 186 mice spanning six visual areas, we aim to better understand population coding in the mouse visual cortex.

Given neural responses from populations of just over one hundred visual cortical neurons, linear classifiers achieve high accuracy in two decoding tasks: one with six stimulus classes with complex spatiotemporal features and one with eight drifting grating directions. We found differential decoding accuracy between the primary (VISp), lateral (VISl), anterolateral (VISal), anteromedial (VISam), posteromedial (VISpm), and rostrolateral (VISrl) visual areas, which implies differential information representation in these visual areas. We also found differences between populations from different cortical depths, with superficial layer populations containing more information than those from deeper layers. Moreover, we found evidence that directional tuning in individual neurons does not necessarily predict the population decoding accuracy suggesting distributed representation of information. These results reveal novel evidence about the cortical organization of visual information processing.

## Materials & Methods

### Dataset

We analyzed data from the Allen Brain Observatory, downloaded on July 3, 2017 using the AllenSDK version 0.13.2. A full description of the Allen Brain Observatory’s data collection methodology is available in their Visual Coding Overview and Visual Stimuli technical whitepapers Allen Institute for Brain Science (2017). In brief, the Allen Brain Observatory recorded *in vivo* two-photon calcium imaging data at 30 Hz over a 400 µm field of view at a resolution of 512 × 512 pixels. We use data from 186 mice of the 216 mice imaged by the Allen Brain Observatory.

Recent studies have identified aberrant cortical activity in GCaMP6-expressing transgenic mouse lines, particularly in Emx1-Cre, a line included in Allen Brain Observatory dataset Steinmetz et al. (2017). By screening somatosensory cortex epifluorescence movies prior to imaging and analyzing visual cortex two-photon calcium recordings after imaging, the Allen Brain Observatory detected aberrant activity resembling epileptiform interictal events in ten Emx-IRES-Cre mice and seven Rbp4-Cre KL100 mice. Data recorded from these seventeen aberrant mice were excluded from our analysis. In addition, data from twelve mice were discarded due to the recording of fewer than ten common neurons across three visual stimulus sessions. Lastly, data from one additional mouse was discarded due to a large number of missing values, resulting in a total of 186 mice with viable data. The sizes (Table S1-S4) and Cre lines (Table S5 and S6) of the populations varied among the targeted visual areas and depths.

A set of synthetic and natural stimuli, comprised of (1) drifting gratings, (2) static gratings, (3) locally sparse noise, (4) natural images, (5) natural movies, and (6) spontaneous activity (mean luminance gray), were displayed on an ASUS PA248Q LCD monitor at a resolution of 1920 × 1200 pixels. Spherical warping was applied to all stimuli to account for the close viewing angle. The monitor was positioned 15 cm from the right eye of awake head-fixed mice, spanning 120° by 95° of visual space without accounting for the spherical warping. The stimuli were distributed into three sessions A, B, and C (or C2) which were presented over three days. The natural movie and spontaneous stimuli were presented in all sessions. Drifting gratings were presented in session A, static gratings and natural images in session B, and locally sparse noise in session C/C2. Session types C and C2 both contained the 4-degree locally sparse noise stimulus (16 × 28 array of 4.65° patches). Session C2 also contained the 8-degree locally sparse noise stimulus (8 × 14 array of 9.3° patches), which was discarded from analysis since it was only shown to a subset of mice.

The drifting gratings stimulus was comprised of 40 grating conditions (combinations of one of 8 directions and one of 5 temporal frequencies), presented 15 times each in a random order. Each grating was presented for two seconds, followed by one second of mean luminance gray.

### Pre-processing

The neural signal was quantified as fluorescence fluctuation Δ*F/F*, calculated for each frame as 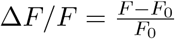, where the baseline *F*_0_ is the mean fluorescence of the preceding 1 second. For each of 186 neural populations, three hours of Δ*F/F* traces were separated into stimulus epochs.

To form samples for the stimulus classification, each epoch was divided into 10 s intervals, of which the final interval was discarded if it was less than 10 s. Neural populations used in the stimulus classification were composed of neurons common across the three imaging sessions A, B, and C (or C2) for each mouse (Tables S1 and S2). For each 10 s interval, the mean fluorescence fluctuation per neuron was calculated and labeled with the corresponding stimulus class.

To form samples for the direction classification, the drifting gratings epoch was divided into 3 s intervals, of which the third second (during which a blank sweep of mean luminance gray was presented) was discarded. Neural populations used in the direction classification were composed of all neurons imaged during session A, and thus were larger than populations used in the stimulus classification (Table S3 and S4). For each 2 s interval, the mean fluorescence fluctuation per neuron was calculated and labeled with the corresponding grating direction.

In both the stimulus and the direction decoding, mean Δ*F/F* for each neuron were z-scored and combined to form the neural feature vectors in ℝ^*n*^ for classification, where *n* is the number of neurons in the population.

### Neural decoding

We used linear classifiers to decode the stimulus classes based on the neural feature vectors. The classifiers were implemented in the Python programming language using the scikit-learn machine learning library version 0.19.0 Pedregosa et al. (2011). Linear support vector machine (SVM) and multinomial logistic regression (MLR) were trained and tested with a nested cross-validation scheme. We principally split the data into training and test sets to form a 5-fold cross-validated prediction.

All results are based on data from both SVM and MLR classification, for which similar results were obtained (Fig. S1). However, we show only show SVM classification results in Figures 2–7 for simplicity.

**Figure 2:**
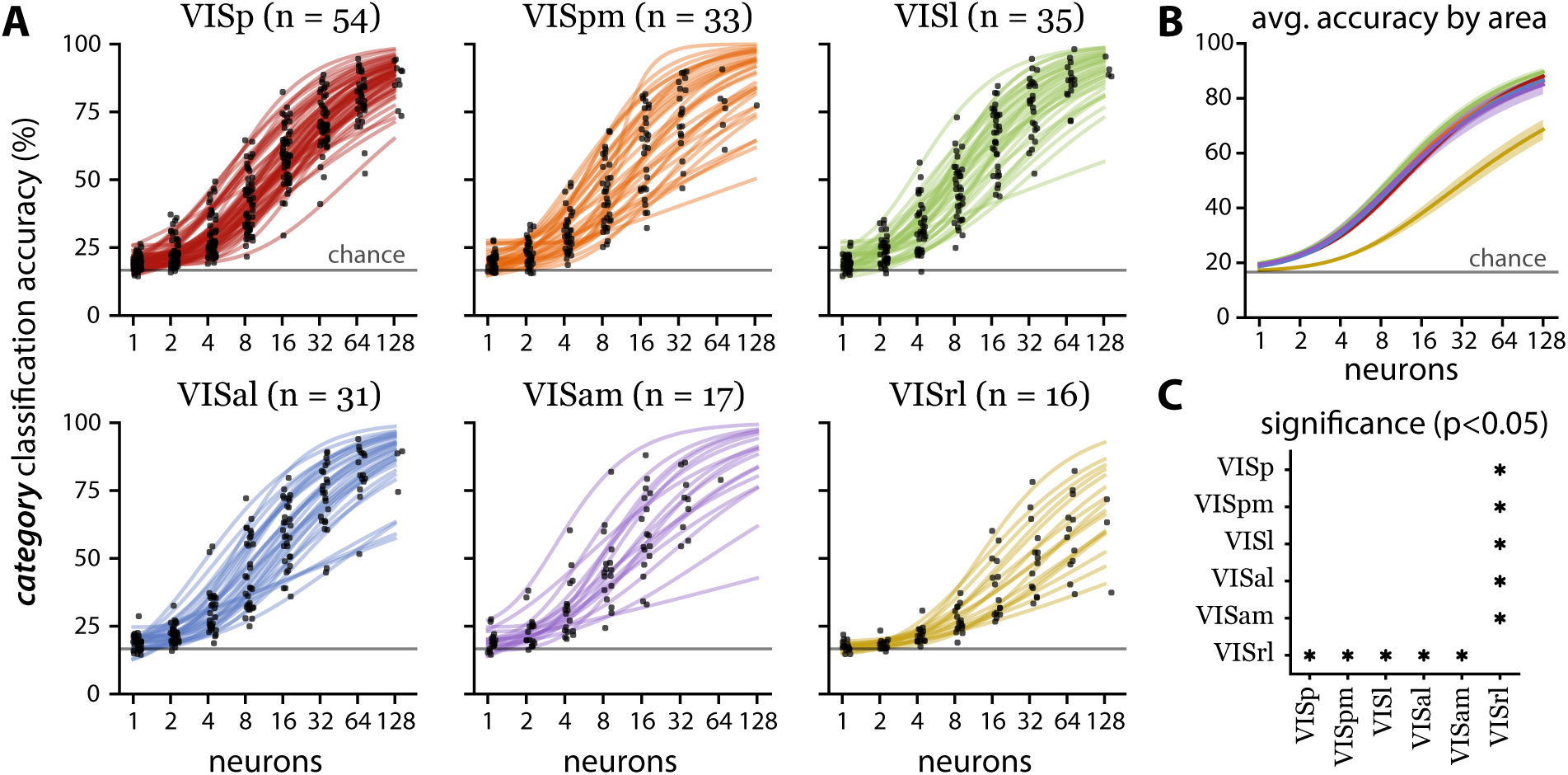
Population decoding performance by visual area for six stimulus classes. (**A**) Stimulus decoding accuracy for individual randomly subsampled populations consisting of 1, 2, 4, 8, 16, 32, 64, and 128 neurons (black dots, jittered for visual clarity) and curve fits (solid lines). The number of populations per area is listed in the titles (*n*). In all visual areas, the majority of small populations (1-4 neurons) outperformed chance level (gray line at 16.67 % accuracy). However, small population performance in VISrl was more concentrated near chance level than all other areas. Individual populations of 128 neurons achieved near-perfect accuracy in all visual areas except VISrl. (**B**) Population averaged accuracy by visual area (solid lines) with standard error (shaded regions). The line colors correspond to the visual area indicated by the line colors in **A**. (**C**) Statistically significant (p *<* 0.05) pairwise comparisons of decoding accuracy at 128 neurons between the six visual areas using Tukey’s test. VISrl underperforms all other visual areas.

**Figure 3:**
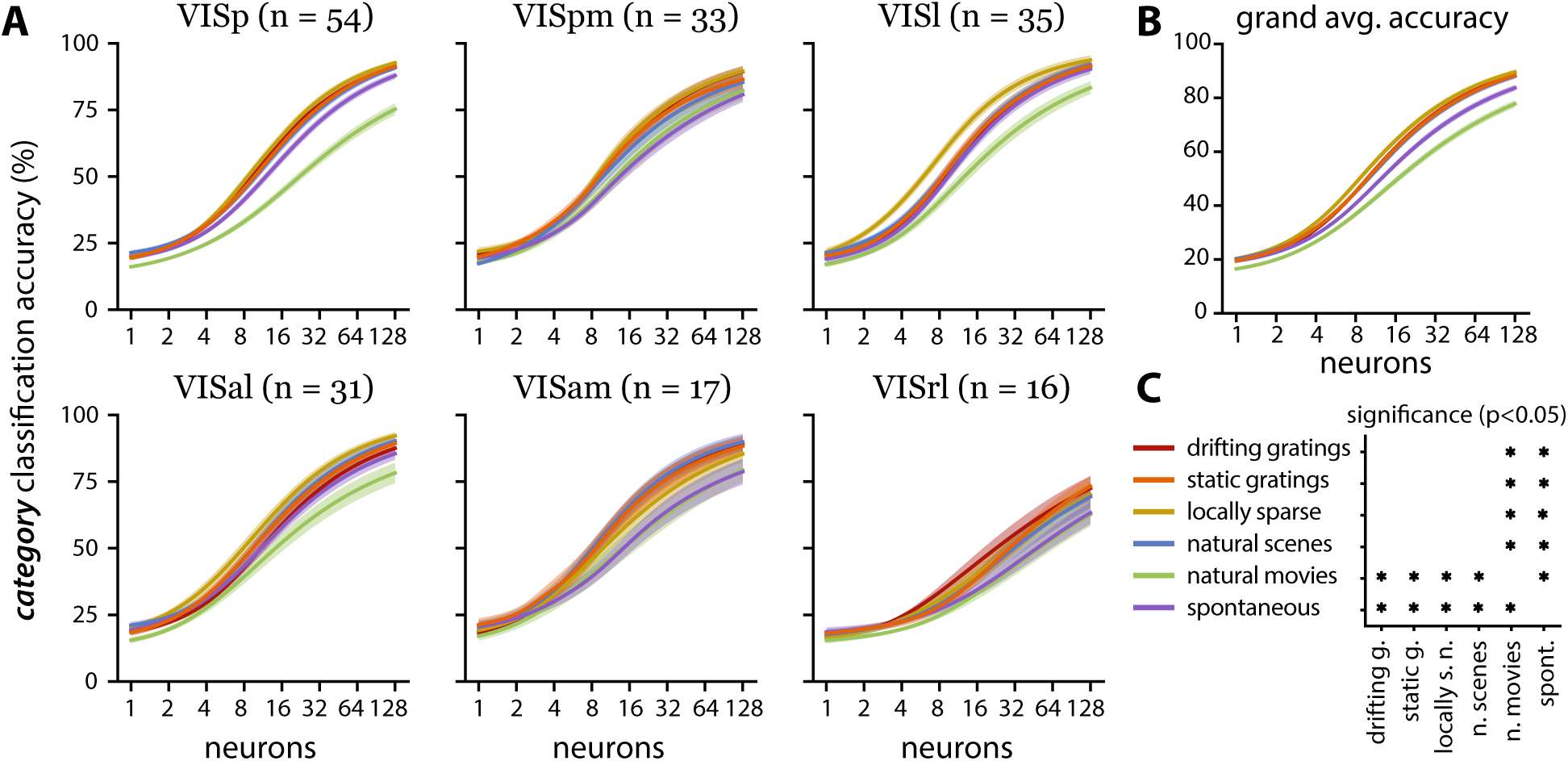
Stimulus-specific population decoding. (**A**) Visual area averaged stimulus-specific decoding accuracy similar to Fig. 2. The line color corresponds to the stimulus indicated by the legend in **C**. High-performing areas with similar overall decoding accuracy show differential accuracy in predicting specific stimuli. (**B**) Stimulus-specific decoding accuracy averaged across all populations. There is differential accuracy in decoding the specific stimulus classes, with some being harder to decode than others. (**C**) Statistical significance map (same convention as Fig. 2C). The natural movies are significantly more difficult to decode than all other stimuli, and the locally sparse noise is significantly more difficult to decode than all others except the natural movies.

**Figure 4:**
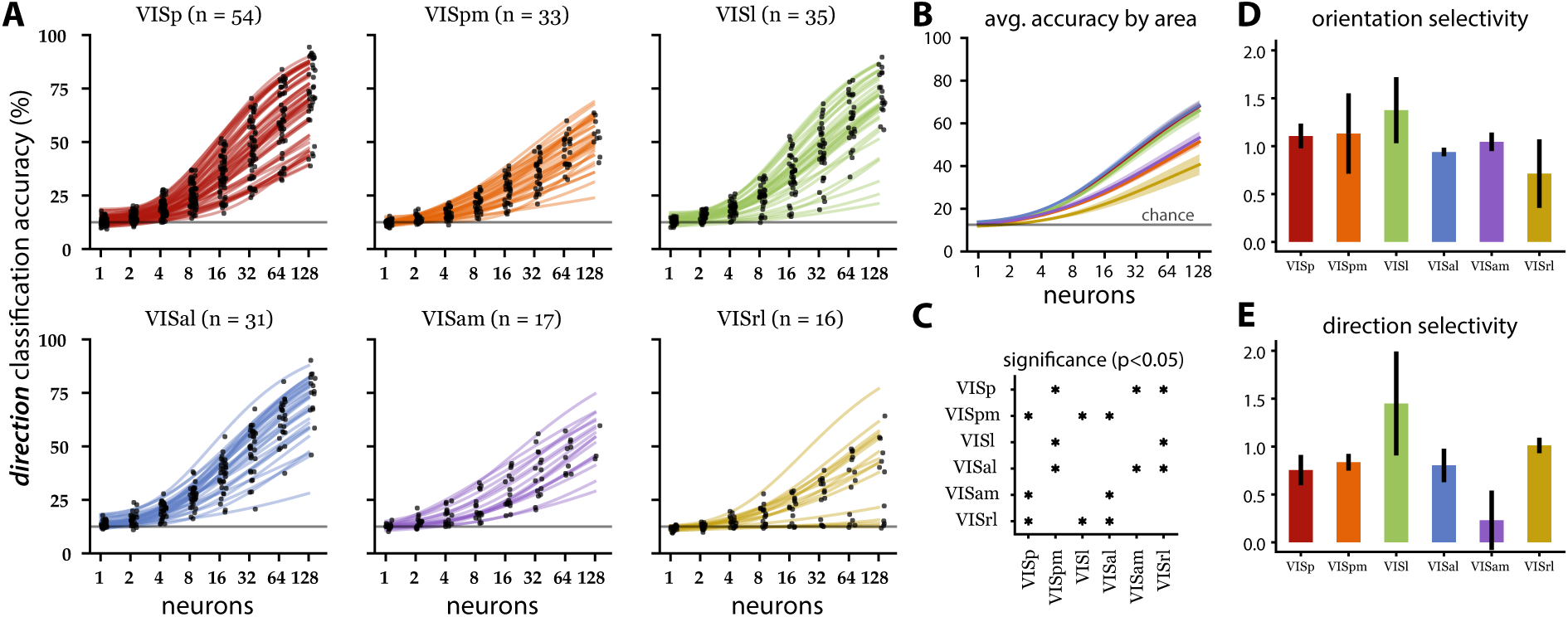
Population decoding of directions for the drifting grating epoch. (**A**) Direction decoding accuracy (same conventions as Fig. 2). Note that in VISrl, small populations (1-2 neurons) performed closer to chance level (gray line at 12.5 % accuracy) than the same sized populations in other areas. (**B**) Population averaged accuracy by visual area (solid lines) with standard error (shaded regions). The line colors correspond to the visual area indicated by the line colors in **A**. (**C**) Statistical significance map (same convention as Fig. 2C). Three high-performing, areas (VISp, VISal, VISl) showing similar performance are anatomically adjacent. Similarly, two of three low-performing areas (VISpm and VISam) showing similar performance are anatomically adjacent. (**D,E**) Mean orientation (D) and direction (E) selectivity index (with SEM) per area (see Methods).

**Figure 5:**
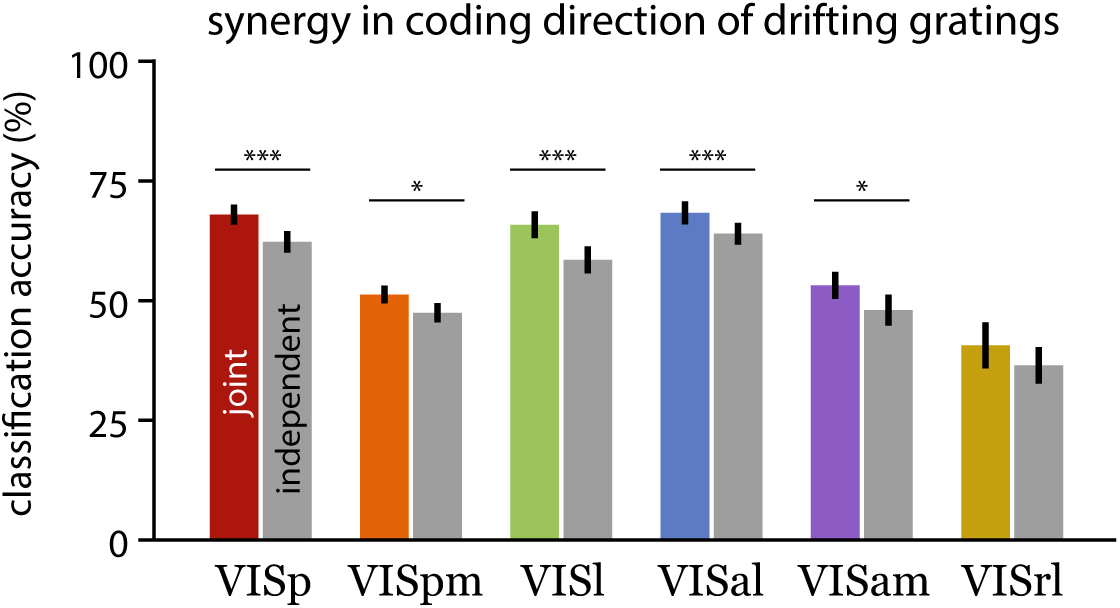
Neural population is synergistically encoding directional information. Accuracy of correlation blind decoder (gray bars; independent decoder) is compared to the joint decoder (same value as in Fig. 4B) for the population size of 128 neurons. Statistical significance indicated by paired t-test.

**Figure 6:**
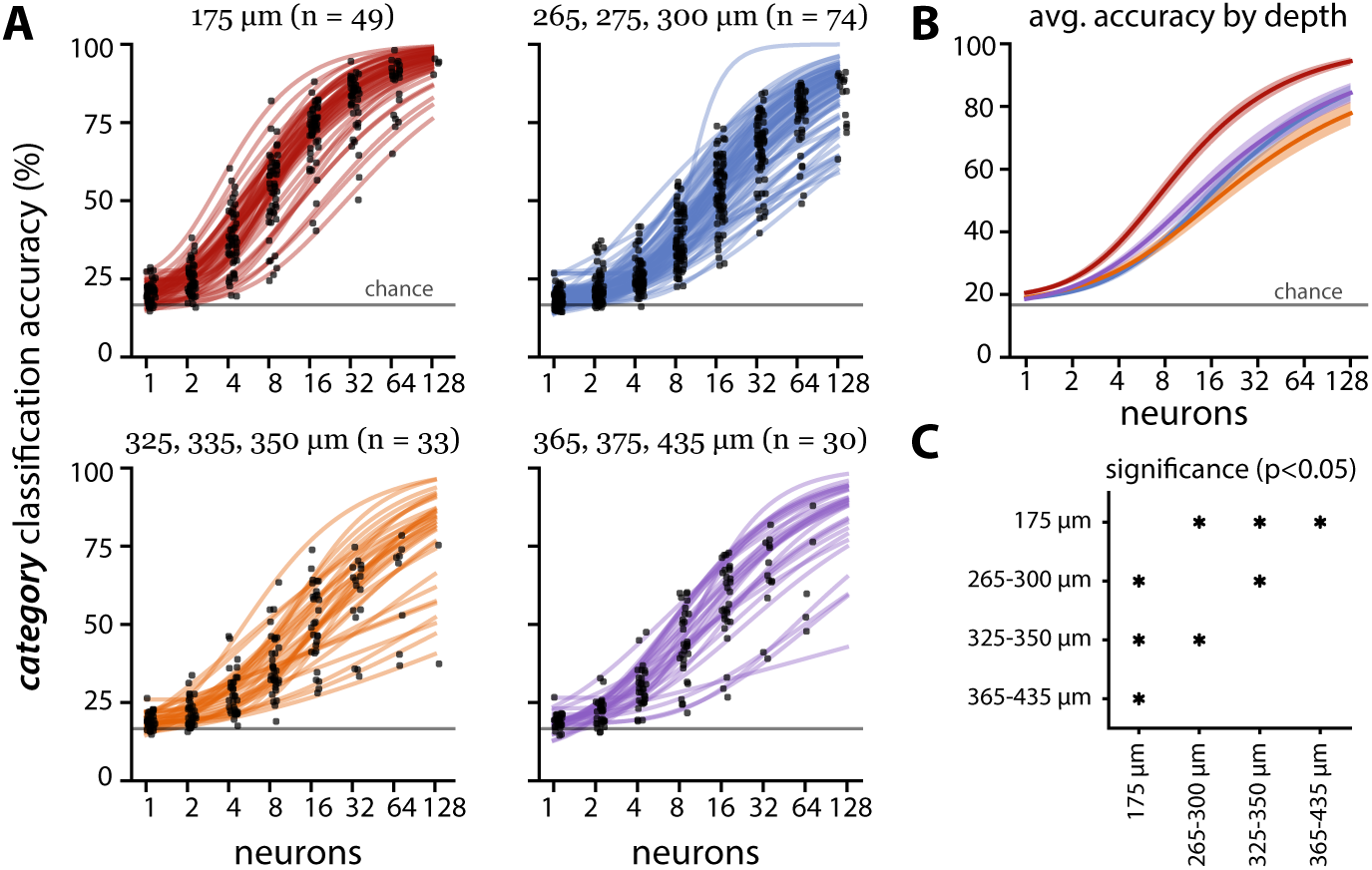
Population decoding performance by recording depth for six stimulus classes. (same conventions as Fig. 2). On average, small populations (1-2 neurons) performed better than chance level performance (gray line at 16.67% accuracy). The 325–350 µm group significantly underperforms two shallower groups (175 µm and 265–300 µm).

**Figure 7:**
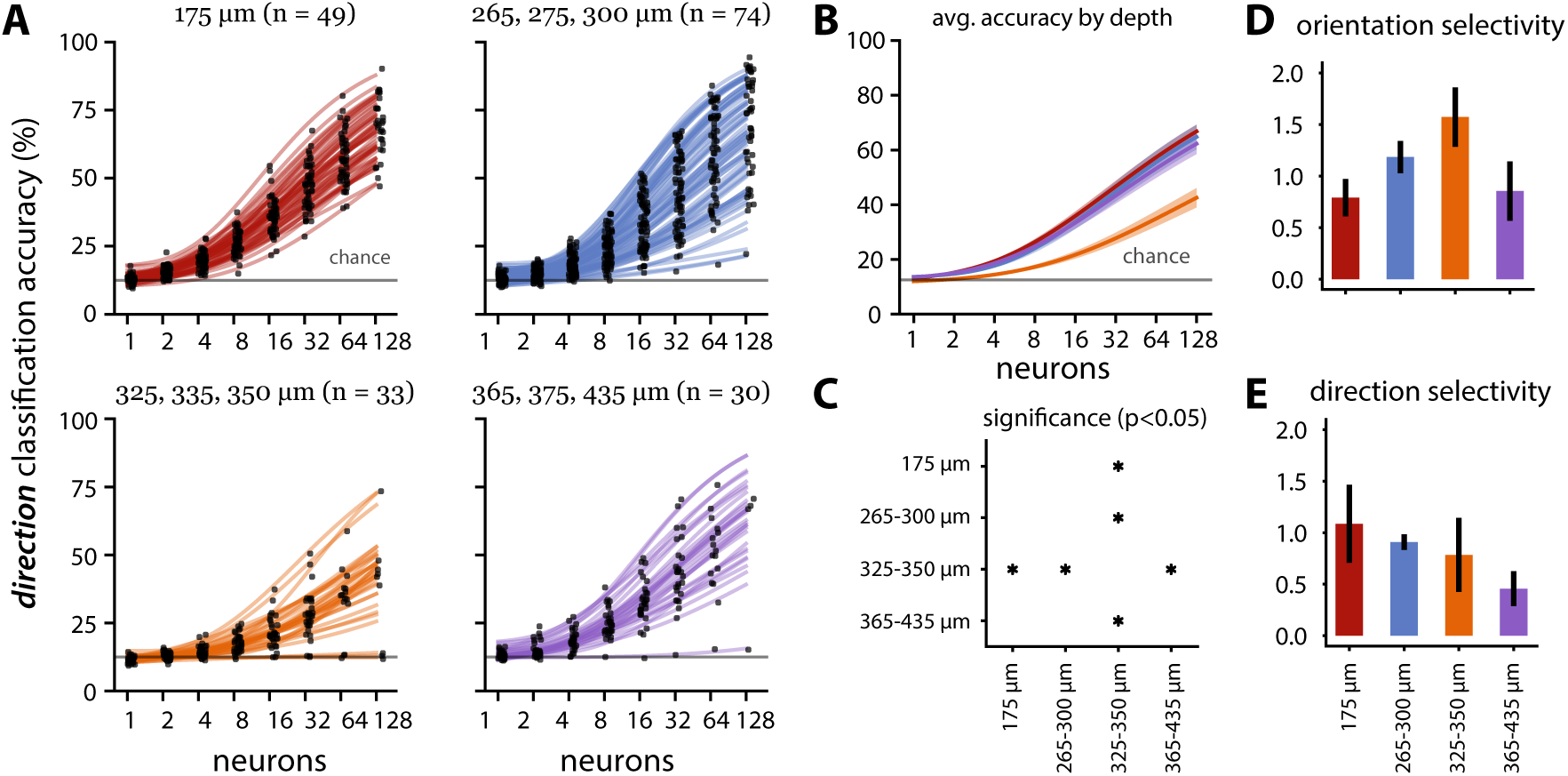
Population decoding performance by imaging depth for eight drifting grating directions. (same conventions as Fig. 4). On average, small populations (1-2 neurons) in the three high performing depth groups (175 µm, 265–300 µm, and 365–435 µm) outperformed chance level (gray line at 12.5% accuracy), while small populations in the low-performing 325–350 µm group performed at chance level.

Due to the different duration of stimulus presentations, the stimulus classes had unbalanced numbers of samples. To build balanced training sets, we subsampled (without replacement) an equal number of responses from each class. The size of these subsamples was equal to 80% of the smallest class (spontaneous activity; 20 minutes out of total 177 or 156 minutes of recording used in samples, depending on if the mouse was shown C or C2). The test sets consisted of the remaining samples, and were kept unbalanced.

The direction classes used in the direction decoding were evenly distributed throughout the stimulus presentation. The direction samples were randomly split intro training (80%) and test (20%) sets for all classification. The training set was assumed to be balanced due to the even distribution of classes throughout data collection.

Both classifiers were regularized using Additive *£*_2_-regularizer of the form 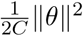. The regularization constant was optimized through a nested cross-validation within the first training set where the best *C* = {10^−2^, 10^−1^, 1, 10, 10^2^, 10^3^, ∞} that yielded the highest accuracy was chosen.

### Subsampled population

To investigate the scaling of decoding performance as a function of population size, we made random subsamples (without replacement) of different sizes *n* = {2^0^, 2^1^, 2^2^, *…*} up to the number of neurons available for each mouse. We repeated the procedure 10 times to form 10 resampled subpopulations. We report accuracy values averaged over the ten resampled datasets. The statistics of population sizes by group or decoding task can be found in Table S1–S4.

To investigate the information carried by the joint population activity, we trained “correlation-blind” decoders with the same procedure but on a shuffled dataset where the joint structure was approximately independent. To generate the shuffled data, we randomly permuted the trials corresponding to the same target for each neuron.

### Accuracy curve fitting

To extrapolate the accuracy as a function of population size, we used the following generalized logistic function:

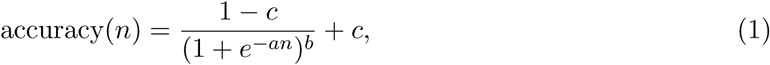

with 3 parameters {*a, b, c*} with constraints *a* ≥ 0, *c* ≥ 0 and *b* ∈ [0, 1]. Note that the *c* parameter allows a minimum accuracy expected from chance level performance for small population size. We fit the curve on the average accuracies obtained by subsampling using nonlinear least squares van der Walt et al. (2011).

### Statistical tests

To compare accuracy between cortical areas and imaging depths, we performed Tukey’s test at a 0.05 significance level Tukey (1949). Tukey’s test compares the mean accuracies of every pair with adjustment for multiple comparison. Ten imaging depths (175 µm, 265 µm, 275 µm, 300 µm, 325 µm, 335 µm, 350 µm, 365 µm, 375 µm, 435 µm) were sorted into four groups: 175 µm, 265-300 µm, 325-350 µm, and 365-435 µm. We compared the six visual cortical areas (VISp, VISpm, VISl, VISal, VISam, and VISrl), four imaging depth groups, and six stimulus classes.

### Orientation and Direction Selectivity

The neural activity recorded during the Session A drifting gratings stimulus was used to calculate the orientation selectivity index (OSI) and direction selectivity index (DSI) for each neuron. We obtained OSI and DSI using the Allen SDK Drifting Gratings module,

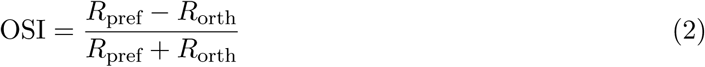

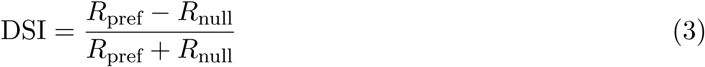

where *R*_pref_ is the mean response to the preferred orientation at the preferred temporal frequency, *R*_orth_ is the mean response to the orthogonal directions, and *R*_null_ is the mean response to the opposite direction. The response was defined as the mean Δ*F/F* during the grating presentation. Each condition was presented 15 times, and responses to all presentations were averaged together. The preferred direction and temporal frequency condition was defined as that grating condition that evoked the largest mean response.

Since Δ*F/F* can be negative, OSI and DSI values can be greater than 1 or even be negative. We excluded values below 0 (663 OSI values and 648 DSI values out of 26186 cells) or above 2 (1871 OSI values and 1561 DSI values) following the Allen Institute guidelines. The full computation methodology for these indices can be found in the Allen Brain Observatory’s Visual Stimuli technical whitepaper Allen Institute for Brain Science (2017). To compare across visual areas, the OSI and DSI of all neurons in each area were averaged together (Fig. 4D,E). To compare across depths, the OSI and DSI of all neurons in each depth were averaged together (Fig. 7D,E).

## Results

### Spatiotemporal structure of stimuli are differentially encoded among visual areas

To investigate differences in information processing between six mouse visual areas, statistical classifiers were fit to discriminate visual categories based on the population activity within each area. Neural activity was monitored through a fluorescent calcium sensor (GCaMP6f) selectively expressed in transgenic mice Allen Institute for Brain Science (2017). Recorded calcium signals were processed and discretized in time to yield feature vectors corresponding to neural activity of the population (see Methods). Mice were shown six types of stimuli which differed in their spatiotemporal structures, ranging from simple spatial structures (such as orientation gratings and sparse pixels) to complex natural scenes (Fig. 1A,B) The stimuli included static images as well as movies with complex long range correlations. A faithful recovery of these visual categories from neural activity reflects the potential information the neural population encodes about the stimuli.

Since the population size was variable across experiments, we compare the rate at which the classification accuracy improves as a function of population size (Fig. 2A). Classification accuracies from small randomly subsampled populations were near chance level, and gradually increased with the population size for all sessions analyzed (Fig. 2A; black dots). We fit a 3-parameter sigmoid function to extrapolate up to 128 neurons for each session (Fig. 2A; see Methods). The averages within each of the six visual areas show similar increasing trends with accuracy approaching 90% for the population size of 128. Five areas (VISal, VISam, VISl, VISp, VISpm) significantly outperformed VISrl (Fig. 2B,C). We used a one-sided t-test with a null hypothesis that secondary areas’ decoding performance is less than that of the primary visual cortex. For both the stimulus category decoding and direction decoding, we failed to reject the null hypothesis at the 0.05 significance level.

We examined the accuracy of decoding specific stimulus categories to further investigate encoding differences across visual areas. On average, natural movie and spontaneous categories were more difficult to decode (Fig. 3B,C). Though similar in overall decoding accuracy, the five high-performing visual areas (VISal, VISam, VISl, VISp, VISpm) show differential pattern in per category accuracy (Fig. 3). We used a one-sided t-test (p-values adjusted for multiple tests) to compare the decoding accuracy of the natural movie stimulus and all other stimulus categories within each visual area. The natural movie category is significantly harder (p *<* 0.001)to decode than other stimuli in populations from the anatomically adjacent VISp, VISl, and VISal (Fig. 3A).

### Area dependent decoding of drifting gratings direction

Local visual orientation information is prevalently encoded in the primary visual cortex Hubel and Wiesel (1959), Priebe (2016). Layer 2/3 neurons in the mouse visual cortex are also sensitive to orientation gratings and their directional motion Marshel et al. (2011). However, mouse primary visual cortex seems to also serve the role of higher order visual function Gavornik and Bear (2014). We investigated if the ability to decode vastly different stimulus categories is related to their capacity to represent orientation and direction. Using the average neural activity in 2-second windows corresponding to the duration of drifting grating presentation, we trained linear classifiers to decode the direction of drifting gratings.

Except for a few VISrl populations, direction decoding was again an increasing function of population size (Fig. 4A). VISrl showed the worst decoding performance at the 128 neuron level, and VISam/VISpm showed intermediate performance, while VISp, VISl and VISal showed comparable population level encoding (Fig. 4B,C).

Surprisingly, the population decoding accuracy showed discrepancies from what is expected from individual neuron’s directional tuning sensitivity. Higher orientation and direction selectivity index (Fig. 4D,E) indicates the stronger representation of these basic visual features, which is highest in VISl followed by VISrl. However, the joint activity decoding showed VISl being on par with VISp and VISal, while the VISrl population was much less informative. This suggests that excitatory neurons in VISp and VISal are more synergistic (a tendency for the population to contain more information than individual neurons Brenner et al. (2017), Latham and Nirenberg (2005)) and that there is relatively more redundancy in the VISl population.

This synergistic population code is corroborated by the general trend of inferior performance of the correlation-blind decoder. The correlation-blind decoder was trained on the trial-shuffled neural data, hence removing the noise correlation. In Fig. 5, for all areas except VISrl there is a significant drop in performance which indicates the joint activity of the population carries extra information.

### Superficial layers are more informative about the spatiotemporal signatures of visual stimuli

In rats, neurons in the superficial layers of V1 are known to have better orientation selectivity and less spontaneous activity Girman et al. (1999), suggesting a laminar organization of visual information processing. To investigate if similar laminar differences exist in mice, we analyzed the decoding accuracy of stimulus classes as a function of recording depth (Fig. 6). There were 6 different Cre lines with specific targets (see Table S6 for full distribution). Since there was little difference across Cre lines, we present the results grouped by depth.

The 325–350 µm depth group (dominated by Nr5a1 Cre line; see Table S6) consistently showed the worst decoding performance across both the stimulus category and direction decoding tasks (Fig. 7). Meanwhile, the most superficial group (imaging depth of 175 µm corresponding to either Cux2 or Emx1 Cre lines, putative layer 2/3) significantly outperformed the deeper populations (Fig. 6), with high decoding performance across all stimulus categories (Fig. S2). However, this superficial layer did not show distinctly superior direction decoding (Fig. 7B). This suggests that the spatiotemporal structure of each visual category extra to the overall orientation information is better represented in the superficial layers. Although there may be worsening of signal-to-noise ratio as the imaging depth increases, both decoding schemes do not show monotonic degradation of performance as a function of depth (Figs. 6 and 7).

The orientation selectivity index (OSI) and DSI showed contrary trends (Fig. 7D,E). Deeper layers had relatively larger OSI but smaller DSI, suggesting the temporal component of the drifting gratings may be better represented in the superficial layers. Despite larger DSI, the 325–350 µm group performed worse than the 365–435 µm group, again an unexpected observation likely due to the spatial organization of the code.

## Discussion

The focus of this study was investigating how stimulus classes and drifting grating directions can be inferred from neural population responses in mouse visual areas. In primates, it has been well established that visual processing occurs through a hierarchical structure, in which the primary visual cortex provides input to secondary visual areas Felleman and Van Essen (1991), Maunsell and Newsome (1987), Orban (2008). The rat visual cortex has also been characterized as having a hierarchical organization Coogan and Burkhalter (1993). Results from this analysis corroborate recent studies which have suggested that this simple hierarchy may also be present in the mouse visual cortex Berezovskii et al. (2011), Wang and Burkhalter (2007). In both decoding tasks, the overall decoding performance of populations from secondary visual areas was equal to or worse than the primary visual cortex (VISp), suggesting that secondary areas do not encode any more information than is encoded by the primary visual cortex. This is supported by findings that the mouse primary visual cortex has a more diverse set of stimulus preferences than secondary areas VISal and VISpm Andermann et al. (2011).

Differences in stimulus-specific decoding performance between populations from different visual areas suggest areal differences in visual information representation. On average, the spontaneous stimulus and the natural movie stimulus are significantly harder to decode than other stimuli, but this trend is not seen in all areas (Fig 3). Anatomically adjacent visual areas display similarities in their stimulus-specific decoding performance. The adjacent anteromedial (VISam) and posteromedial (VISpm) areas showed no difference in performance for specific stimuli. In contrast, in populations from the adjacent primary (VISp), anterolateral (VISal), and lateral (VISl) visual areas, it was significantly harder to decode the natural movie stimulus than other stimuli. These anatomical trends in stimulus-specific decoding may be attributed to specialized input pathways from the primary visual cortex Marshel et al. (2011).

The existence of these information processing streams is further supported by the similar direction decoding performance of anatomically adjacent areas. The same groups emerge in the direction decoding as in the stimulus-specific analysis. The adjacent primary (VISp), anterolateral (VISal) and lateral (VISl) visual areas performed similarly, as did the adjacent anteromedial (VISam) and posteromedial (VISpm) areas. The poor performance of the latter group of visual areas (VISam and VISpm) as well as the rostrolateral (VISrl) visual area suggests a lack of direction sensitive information encoding in the population. We speculate that the relative poor performance of VISrl compared to VISam in the population decoding to be in the distribution of well tuned neurons—VISam had lower single neuron DSI on average, but more heterogeneous distribution.

Marshel et al. (2011) presented drifting bar and drifting grating stimuli to 28 mice and found, based on the mean DSI of each area and the proportion of neurons with a DSI greater than 0.5, that layer 2/3 (130-180 µm below the dura surface) populations in the anterolateral (VISal), rostrolateral (VISrl), and anteromedial (VISam) visual areas were significantly more direction selective than the primary visual cortex (VISp). The results of our population direction decoding analysis (Figs. 4 and 5) of 186 mice are inconsistent with the single neuron findings of Marshel et al. (2011) (note that there were differences in the methods for estimating DSI; see Methods). The direction decoding accuracy of VISam and VISrl populations are significantly lower than that of VISp, suggesting that these populations are less direction selective than those in VISp. Trial shuffled decoding analysis (Fig. 5) showed that synergistic spatial correlations within trial could contribute to such discrepancies Averbeck et al. (2006), Brenner et al. (2017). Furthermore, the similar decoding accuracy of VISal and VISp populations suggests that VISal is not significantly different from VISp in its direction selectivity.

Across all visual areas, individual neurons encode enough attributes of a stimulus in their responses that the majority of small populations outperformed chance level accuracy in the stimulus decoding (chance equal to 16.67 %) as well as in the direction decoding (chance equal to 12.5 %). However, in the direction decoding, individual neurons from VISrl populations and those from the 325–350 µm depth group performed at chance level, suggesting a lower proportion of direction sensitive encoding in these neurons relative to other areas and depths. Neurons in VISam have previously been characterized as extremely robust and selective Marshel et al. (2011). However, our direction decoding analysis shows that decoding accuracy for small VISam populations of 1-4 neurons remains at or close to chance level, suggesting that these neurons are not especially selective. Even with larger VISam populations, direction decoding accuracy remained low relative to other areas.

Despite some discrepancies with recent characterizations of mouse visual areas, this study provides novel evidence of the functional and anatomical organization of the mouse visual cortex. The results corroborate broad trends in visual information processing, supporting the existence of information processing streams and a hierarchical organization in the mouse visual cortex.

## Acknowledgement

We would like to thank Michael Buice for thoughtful discussions. We would also like to thank the Allen Institute for Brain Science for supporting open data and open science. This work was partially supported by the Simons Summer Research Program hosted by Stony Brook University.

## Supplementary material

**Figure S1:**
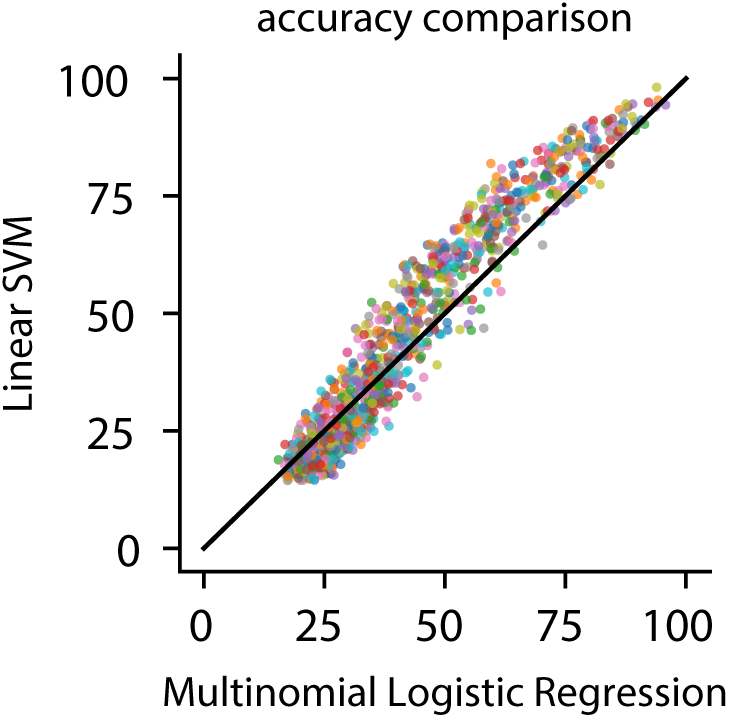
Comparison of results of linear support vector machine (SVM) and multinomial logistic regression (MLR) classification. SVM and MLR classification accuracy for subsampled populations of 1, 2, 4, 8, 16, 32, 64, and 128 neurons are represented by a single point. Similar results were achieved by both classifiers.

**Table S1:**
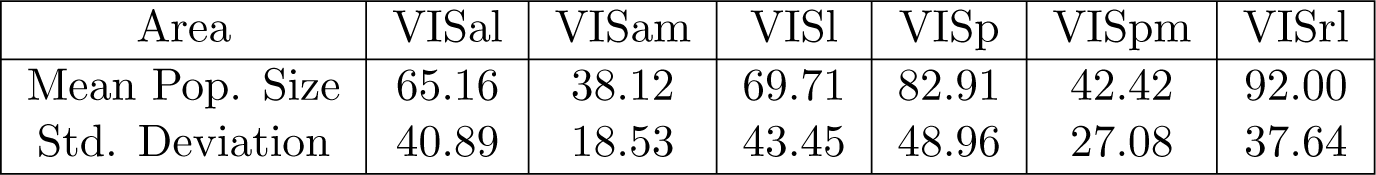
Mean population size with standard deviation by visual area for the stimulus classification. Populations are composed of neurons common across the three imaging sessions A, B, and C.

| Area | VISal | VISam | VISl | VISp | VISpm | VISrl |
| --- | --- | --- | --- | --- | --- | --- |
| Mean Pop. Size | 65.16 | 38.12 | 69.71 | 82.91 | 42.42 | 92.00 |
| Std. Deviation | 40.89 | 18.53 | 43.45 | 48.96 | 27.08 | 37.64 |

**Table S2:** Mean population size with standard deviation by imaging depth group for the stimulus classification. Populations are composed of neurons common across the three imaging sessions A, B, and C.

| Imaging Depth ($\mu\text{m}$) | 175 | 265-300 | 325-350 | 365-435 |
| --- | --- | --- | --- | --- |
| Mean Pop. Size | 70.82 | 84.96 | 46.82 | 38.50 |
| Std. Deviation | 33.74 | 50.14 | 34.20 | 22.62 |

**Figure S2:**
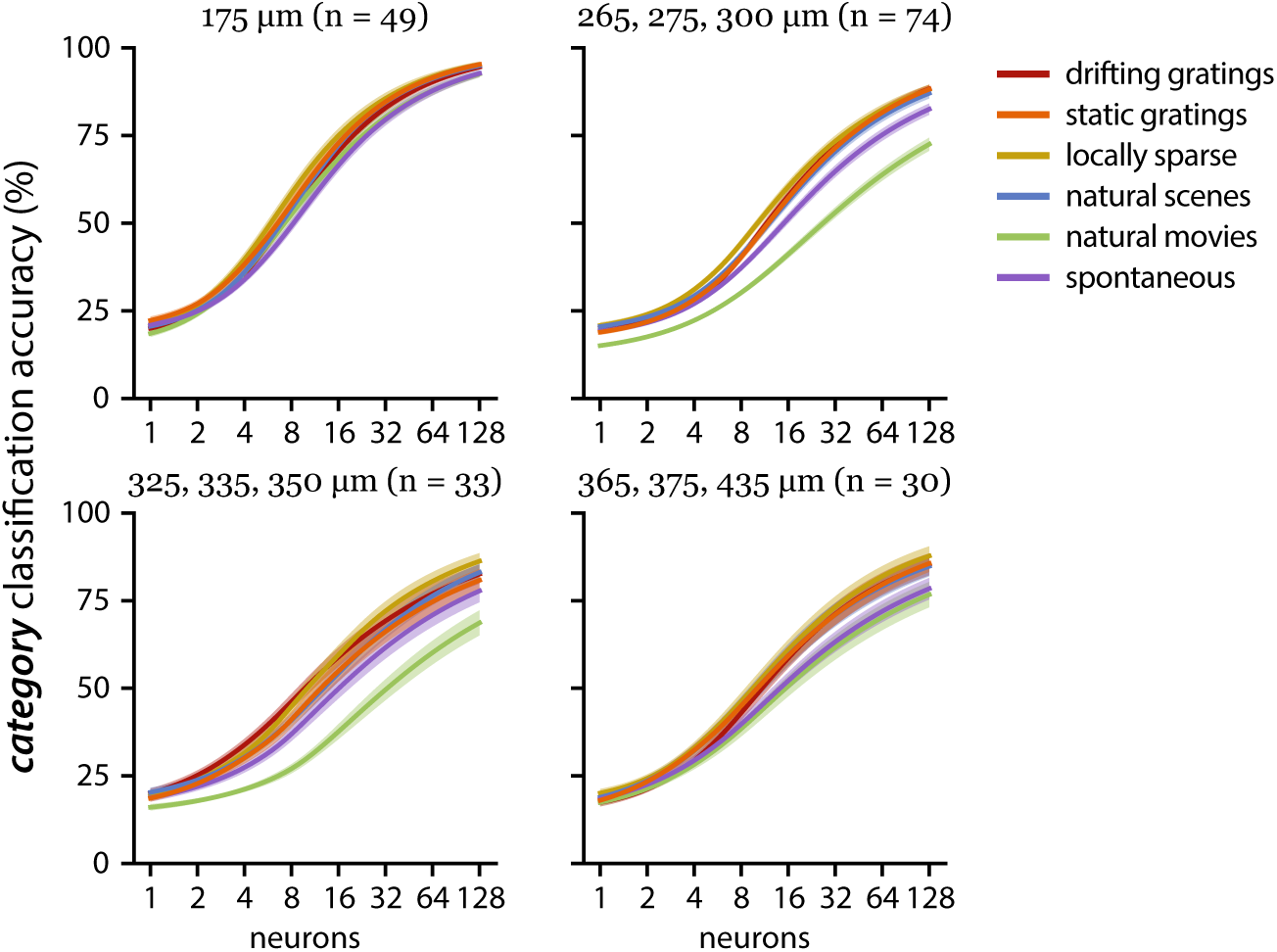
Stimulus-specific decoding performance by imaging depth group. The highest performing depth (175 µm) and a lower performing group (365-435 µm) show uniform accuracy in decoding all six stimuli. In the 265-300 µm and 325-350 µm groups, natural movies are significantly harder to decode than other stimuli.

**Table S3:** Mean population size with standard deviation by visual area for the direction classification. Population sizes are larger for the direction classification than the stimulus classification because the population includes all neurons imaged in Session A.

| Area | VISal | VISam | VISl | VISp | VISpm | VISrl |
| --- | --- | --- | --- | --- | --- | --- |
| Mean Pop. Size | 139.13 | 89.12 | 143.97 | 169.87 | 99.87 | 178.13 |
| Std. Deviation | 79.40 | 56.77 | 85.16 | 84.83 | 62.29 | 68.83 |

**Table S4:** Mean population size with standard deviation by imaging depth group for the direction classification. Population sizes are larger for the direction classification than the stimulus classification because the population includes all neurons imaged in Session A.

| Imaging Depth ($\mu\text{m}$) | 175 | 265-300 | 325-350 | 365-435 |
| --- | --- | --- | --- | --- |
| Mean Pop. Size | 153.49 | 178.15 | 93.27 | 80.13 |
| Std. Deviation | 56.76 | 88.79 | 66.22 | 52.75 |

**Table S5:** Cre line populations in each visual area.

| Area | VISal | VISam | VISl | VISp | VISpm | VISrl |
| --- | --- | --- | --- | --- | --- | --- |
| Cux2-CreERT2 | 11 | 5 | 13 | 16 | 13 | 3 |
| Emx1-IRES-Cre | 7 | 2 | 6 | 7 | 4 | 6 |
| Nr5a1-Cre | 4 | 1 | 5 | 9 | 6 | 4 |
| Rbp4-Cre_KL100 | 3 | 4 | 5 | 5 | 5 | 1 |
| Rorb-IRES2-Cre | 6 | 5 | 6 | 8 | 5 | 2 |
| Scnn1a-Tg3-Cre | 0 | 2 | 0 | 9 | 0 | 0 |

**Table S6:** Cre line populations in each depth group.

| Imaging Depth ($\mu\text{m}$) | 175 | 265-300 | 325-350 | 365-435 |
| --- | --- | --- | --- | --- |
| Cux2-CreERT2 | 34 | 27 | 0 | 0 |
| Emx1-IRES-Cre | 15 | 10 | 0 | 7 |
| Nr5a1-Cre | 0 | 3 | 26 | 0 |
| Rbp4-Cre_KL100 | 0 | 0 | 0 | 23 |
| Rorb-IRES2-Cre | 0 | 32 | 0 | 0 |
| Scnn1a-Tg3-Cre | 0 | 2 | 7 | 0 |

